# Collective mechanical adaptation of honeybee swarms

**DOI:** 10.1101/188953

**Authors:** O. Peleg, J.M. Peters, M.K. Salcedo, L. Mahadevan

## Abstract

Honeybee *Apis mellifera* swarms form clusters made solely of bees attached to each other, forming pendant structures on tree branches (*1*). These clusters can be hundreds of times the size of a single organism. How these structures are stably maintained under the influence of static gravity and dynamic stimuli (e.g. wind) is unknown. To address this, we created pendant conical clusters attached to a board that was shaken with varying amplitude, frequency and total duration. Our observations show that horizontally shaken clusters spread out to form wider, flatter cones, i.e. the cluster adapts to the dynamic loading conditions, but in a reversible manner - when the loading is removed, the cluster recovers its original shape, slowly. Measuring the response of a cluster to a sharp pendular excitation before and after it adapted shows that the flattened cones deform less and relax faster than the elongated ones, i.e. they are more stable mechanically. We use particle-based simulations of a passive assemblage to suggest a behavioral hypothesis that individual bees respond to local variations in strain. This behavioral response improves the collective stability of the cluster as a whole at the expense of increasing the average mechanical burden experienced by the individual. Simulations using this rule explain our observations of adaptation to horizontal shaking. The simulations also suggest that vertical shaking will not lead to significant differential strains and thus no adaptation. To test this, we shake the cluster vertically and find that indeed there is no response to this stimulus. Altogether, our results show how an active, functional super–organism structure can respond adaptively to dynamic mechanical loading by changing its morphology to achieve better load sharing.

---

Collective dynamics allow super–organisms to function in ways that a single organism cannot, by virtue of their emergent size, shape, physiology and behavior (*2*). Classic examples include the physiological and behavioral strategies seen in social insects, e.g. ants that link their bodies to form rafts to survive floods (*3–6*), assemble pulling chains to move food items (*8*), form bivouac (*7*), as well as bridges and ladders to traverse rough terrain (*9*). Similarly, huddling groups of “daddy longlegs”, emperor penguins cluster together for thermoregulation purposes (*10*), etc. While much is known about the static forms which are seen in such situations, the stability of these forms to dynamic perturbation, and their global response in the context of adaptation is much less understood.

European honeybees, *Apis mellifera*, show many of these collective behaviors during their life cycle (*1*). For example, during the reproductive cycle of bees: colonies reproduce through colony fission, a process in which a subset of the colony’s workers and a queen leave the hive, separate from the parent colony, and form a cluster on a nearby tree branch (*1*). In these swarm clusters (which we will refer to as clusters), the bees adhere to each other and form a large structure made of ~ 10,000 individuals and can be hundreds of times the size of a single organism (Fig. 1A). Generally, this hanging mass of adhered bees takes on the shape of an inverted pendant cone, however, the resultant shape is also influenced by the surface to which the cluster is clinging to (see two different examples in Fig. 1A). The cluster can stay in place for several days as scout bees search the surrounding area for suitable nest sites (*1*).

**Figure 1:**
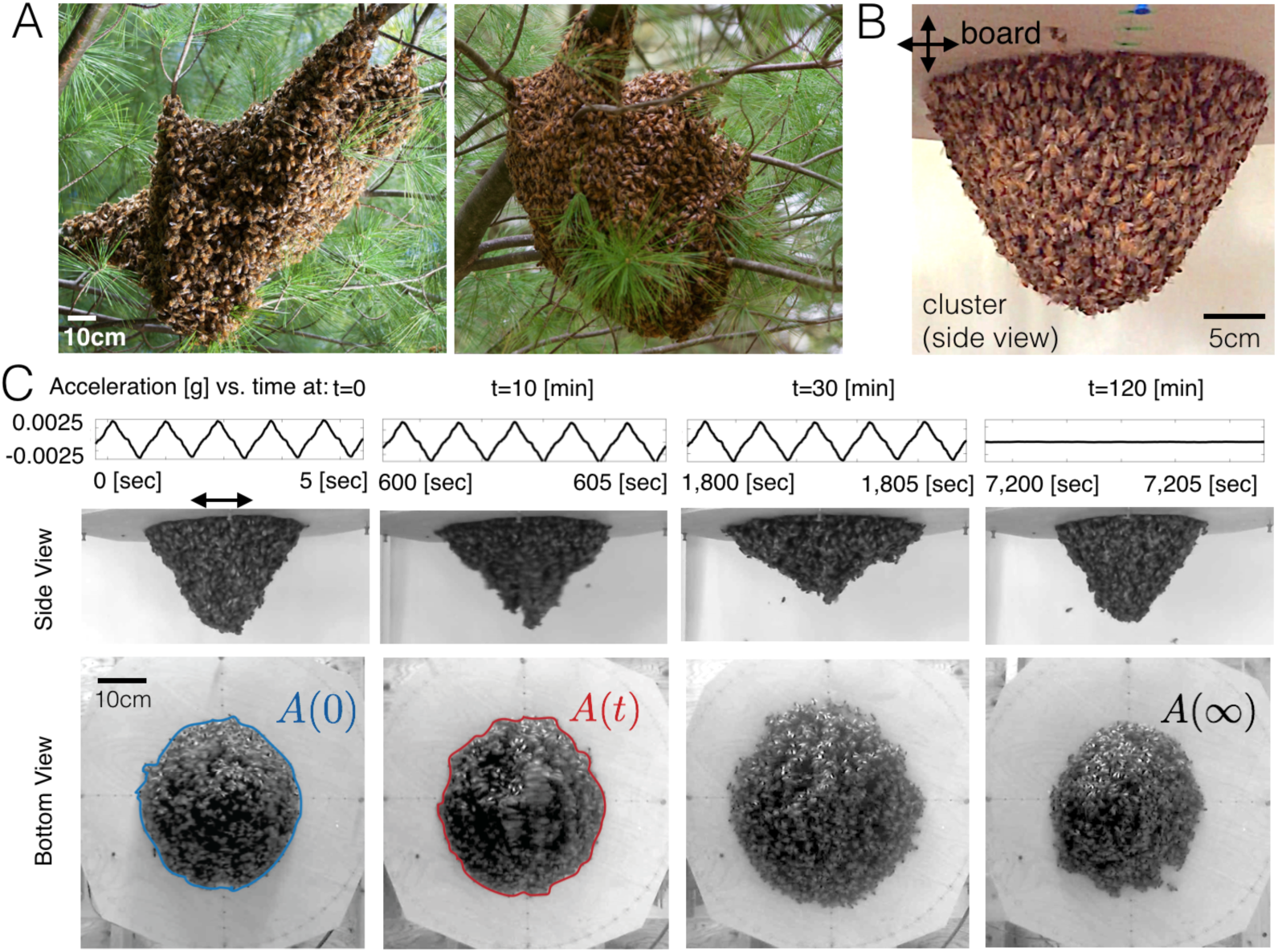
A mechanically adaptive honeybee cluster. A) Bee clusters on a tree branch. B) Experimental setup consists of a motor driving a wooden board, on which a cluster of bees grips a roughly circular contact area. The motor can produce periodic movement in the horizontal or vertical axis at different frequencies and amplitudes. See Fig. S1 for full setup. C) The top panel shows the acceleration of the board vs. time. The middle and bottom panels show how the bee cluster adapts its shape dynamically: Elongated cluster at t=0 (left column), spread-out cluster after horizontal shaking for 10 minutes and 30 minutes (middle columns), and elongated cluster after relaxation (right column); side and bottom views. The contact area before and after shaking is highlighted in blue and red, respectively.

The colony is never more exposed to the environment than during this stage and these clusters show several striking behaviors to cope with the fluctuating thermal and mechanical environment. For instance, clusters tune their density and surface area to volume ratio to maintain a near constant core temperature despite large fluctuations in the ambient temperature (*11–13*). Furthermore, at high temperatures, the swarm expands and forms channels which are presumed to aid in air circulation (*14*). Additionally, in response to rain, bees at the surface arrange themselves to form “shingles”, shedding moisture efficiently from the surface of the cluster (*15*). Similarly, the cluster is mechanically stable; while it sways from side to side in the wind (e.g. see Movie S1), it could be catastrophic if the cluster breaks (when a critical load appears) as the bees would lose the ability to minimize surface area to prevent hypothermia, while still being mechanically stable. However, the mechanism by which a multitude of bees work together to create and maintain a stable structure that handles both static gravity and dynamic shaking stimuli (e.g. wind, predators), remains elusive. To understand this, we develop an experimental setup to quantify the response of a honeybee cluster to mechanical shaking over short and long times.

To prepare a cluster, we attach a caged queen (see SI Sec. A) to a board and allowed a cluster to form around her (Fig. 1B). The bees at the base grip onto an area that is roughly circular. The board is controlled by a motor that can produce movement in the horizontal direction at different frequencies (0.5Hz – 5Hz) and accelerations (ranged 0 – 0.1g). We apply both discontinuous shaking in which the acceleration is kept constant and the frequency is modified, and vice versa, continuous shaking in which the frequency is kept constant and the acceleration is modified (see Fig. S2).

For the case of horizontal shaking (for both discontinuous and continuous), the tall conical cluster swings to and fro in a pendular mode (one of the lowest energy modes of motion, see SI Sec. C), with a typical frequency of ~ 1Hz. However, over longer durations (i.e. minutes), the bees adapt by spreading themselves into a flatter conical form (Fig. 1B,C,D, Movie S2), while their total number remain constant (measured by the total weight of the cluster). The final shape flattens as the shaking continues for longer, or as frequency and acceleration of shaking increases. For the discontinuous shaking, when we plot the relative extent of spreading (scaled by a constant) as measured by *A*(*t*)/*A*(0) for all different frequencies, as a function of number of shakes, the data collapses onto a single curve (Fig. 2A). This suggests that the cluster response scales with both the number and magnitude of shakes, but over much longer time scales than an individual event. The nature of this response is independent of the type of stimulus: when the shaking signal is continuous, we see a similar response (Fig. 2B). The graded adaptive response that scales with the number of shakes and is a function of applied displacements and frequencies, and the absence of any adaptation to very low frequencies and amplitudes (orange curves in Fig. 2B) suggests that there is a critical relative displacement (i.e. a threshold mechanical strain) needed to trigger this adaptation. Once the shaking stops, the cluster returns to its original elongated cone configuration over a period of 30 – 120 minutes, a time that is much larger than the time for the cluster to flatten. This reversible cluster shape change to dynamic loading might be a functional adaptation that increases the mechanical stability of a flattened cluster relative to an elongated one.

**Figure 2:**
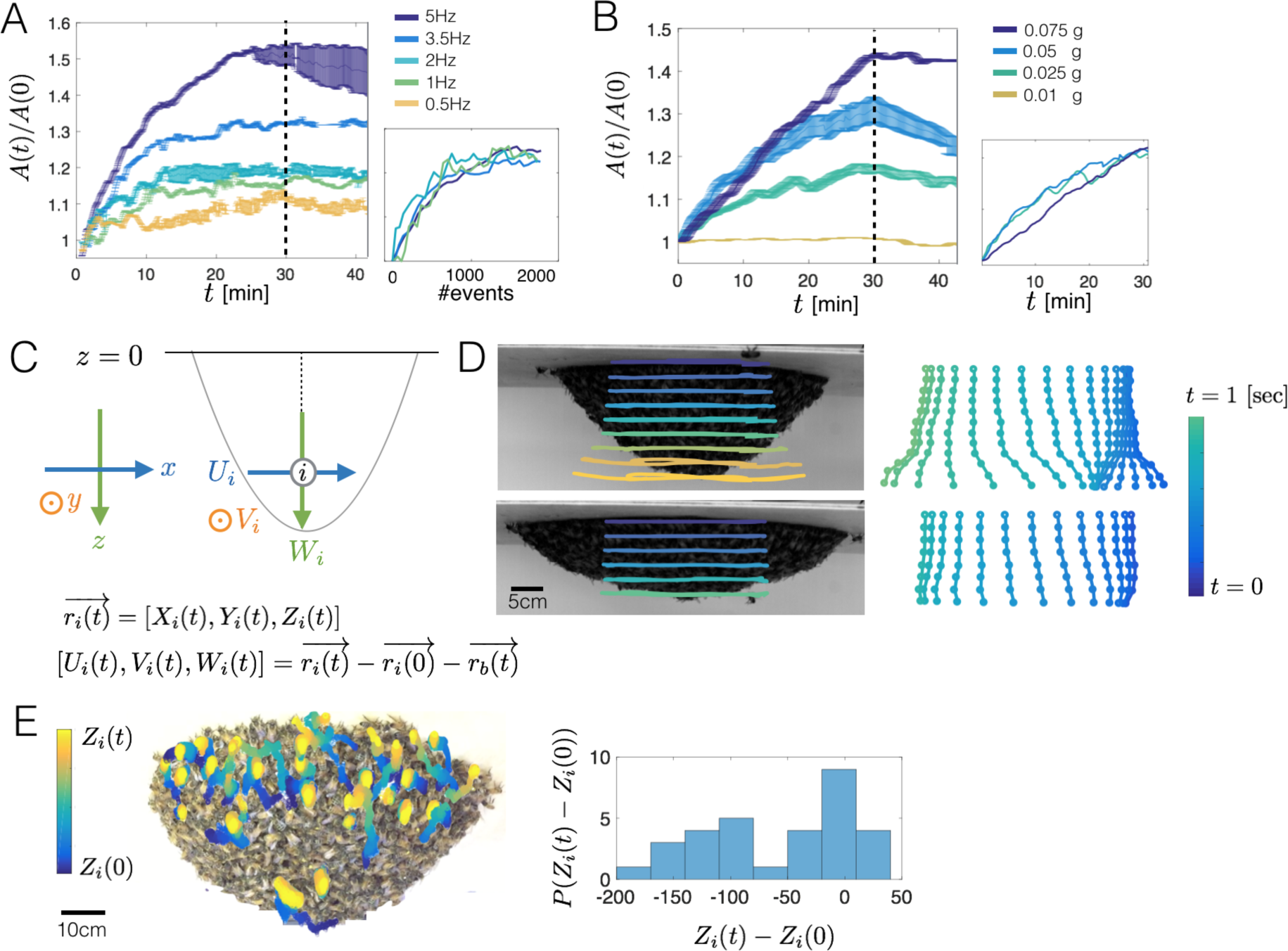
Quantifying adaptive response of the cluster: For all shaking frequencies, the base contact area of the cluster increases monotonically until a plateau is reached. Once shaking ceases, the cluster responds by gradually reverting to its original shape by increasing its contact area, but at a much slower rate. A) Ratio of the contact area of the base of the cluster divided by its original area *A*(*t*)/*A*(0) as a function of time, for the discontinuous case. Colors represent results for different frequencies of periodic shaking. Inset shows that the scaled base area collapses onto a master curve when plotted vs. number of shaking events. Error bars correspond to standard deviation of three individual trials. B) *A*(*t*)/*A*(0) for continuous shaking shows the same qualitative behavior; note that when the acceleration is very small (0.01g), there is no response, i.e. there is a critical threshold of forcing below which the bees do not respond. C) Coordinate systems of the lab-frame and the displacement coordinates of the individual bees. D) Deformation of an elongated cluster before shaking began (*t* = 0, top) and a flattened cluster after shaking (*t* = 30 [sec], bottom) shows that displacement at the tip of the cluster is largest. On the right: time snapshots of a string of bees along the center of the cluster (See Movie S3). E) Trajectories of individual bees during 5 minutes of horizontal shaking show that when the cluster spreads out, surface bees move upwards. Color code represents time: the trajectory starts with blue and ends with yellow. Inset: probability distribution function of vertical displacement, showing a net upward trend.

To explore this suggestion quantitatively, we first define a lab-fixed coordinate system with axes as shown in Fig. 2C, with respect to which the board is at 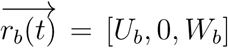 the position of a bee *i* is defined as 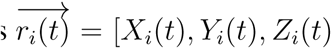, and its displacement is defined as 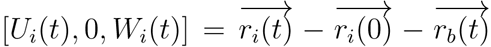. This allows us to track individual bees (*18*) along the surface of the cluster along the centerline *X*_*i*_(0) = 0 (Fig. 2D, Movie S3), over a period of oscillation. Comparing trajectories of bees in an elongated cluster and a flat cluster, i.e. before and after shaking, show that relative displacement between the bees at the cluster tip and bees at the base is significantly larger for an elongated cluster. Snapshots of tracked bees highlight the decoupling of movement of the tip and base of the cluster, i.e. local deformations such as normal and shear strains are reduced in the mechanically adapted state corresponding to a spread cluster. A similar trend is observed when the cluster is subjected to a single sharp shake (see signal at Fig. S2C), as shown in Movie S4. These measurements confirm that the adapted flattened structure is indeed more mechanically stable in the presence of dynamic horizontal loads.

Spreading of the cluster is a collective process, begging the question of how this collective spreading behavior is achieved. To study this, we tracked bees on the surface of the cluster during the process of adaptive spreading, particularly at the early stages. In Fig. 2E, Movie S5 we show how bees move from the tip-regions that are subject to large relative displacements towards the base-regions that are subject to small relative displacements, suggesting that the relative displacement *U*_*i*_, may be a driver of shape adaptation.

But what measure of the relative displacements might the bees be responding to? To understand this, we note that the fundamental modes of a pendant elastic cone are similar to those of a pendulum swinging from side to side, and a spring bouncing up and down, and their frequencies monotonically increase as a function of the aspect ratio of the cluster (Fig. S3) (see SI Sec. C for details). To quantify the deviations from this simple picture due to the particulate nature of the assemblage, we turn to a computational model of the passive dynamics of a cluster and explore the role of shape on a pendant mechanical assemblage of passive particles used to mimic bees. We model each bee in the cluster as a spherical particle which experiences three forces: a gravitational force, an attractive force between neighboring particles, and a force that prevents inter-particle penetration (see SI Sec. C for further details). The bees at the base are assumed to be strongly attached to the supporting board, and those on the surface are assumed to be free. To study the passive response of the entire system, the board is oscillated at different frequencies and amplitudes, while we follow the displacement of individual 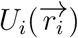, as well as the relative displacement between neighboring bees 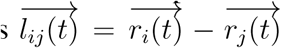 (Fig. 3A). Decomposing the vector 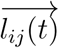 into its magnitude and direction allows us to define two local deformation measures associated with the local normal strain and shear strain. The local dynamic normal strain associated with a particle (bee) i relative to its extension at *t* = 0 is defined as 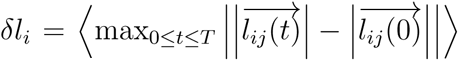 where *T* is the duration from the onset of the applied mechanical shaking until the swarm recovers its steady state configuration, and the brackets <> represent average over all bees *j* that are connected to bee *i*. The local shear strain is calculated from the changes in the angle 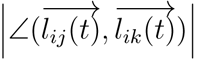 between 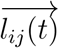 and 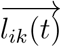 connecting bees *i* and *j*, and bees *i* and *k*, respectively, with the shear strain, *δθ*_*i*_ defined as 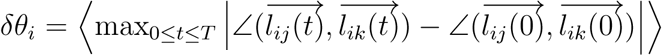 where the brackets <> represent average over all pair of bees *j* – *k* that are connected to bee *i*.

**Figure 3:**
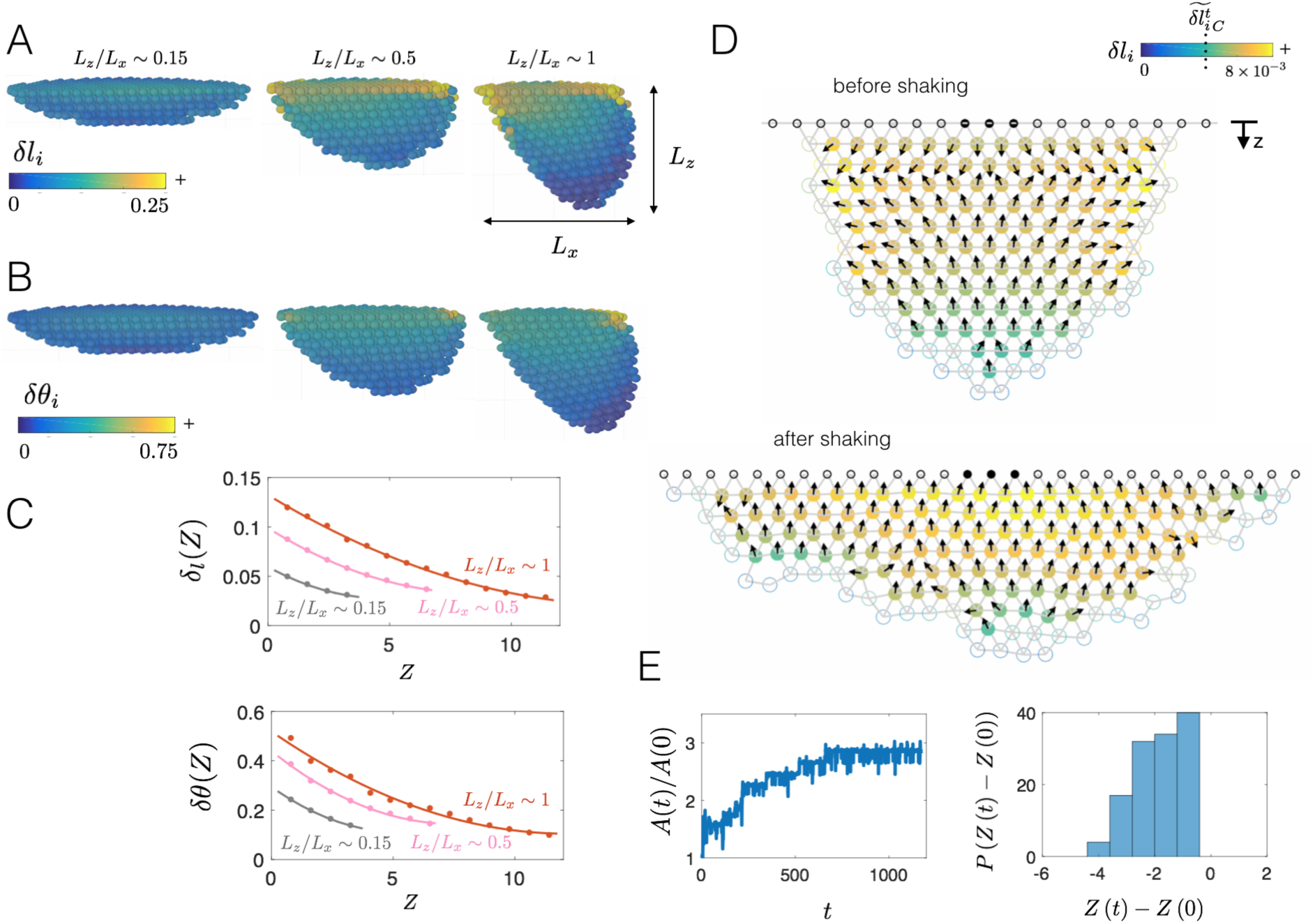
Computational model of mechanical adaptation: A cluster is modeled using particles that are linked via springs in a simple 2d triangular lattice. A) Clusters of different aspect ratios (*L*_*z*_/*L*_*x*_), shown at the extreme of a period of horizontal oscillation. Colors represent the local normal strain of each honeybee *δl*_*i*_, as defined in the text. Elongated clusters (on the right) experience a larger deformation at the tip of the cluster, while flattened clusters (on the left) experience much less deformation. B) For the same state as in A), we also show the maximum shear strain, *δθ*_*i*_. C) Plots of the mean normal and shear strain (*δl*(*Z*) and *δθ*(*Z*)) as a function of the distance from the base, *Z*, and aspect ratio *L*_*z*_/*L*_*x*_. We see that the maximum magnitude of the strains decreases as the cluster becomes flattened. D) When we impose a behavioral rule that allows the bees to sense the strains around them and move in the direction of increasing strain when the magnitude crosses a threshold 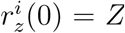, this leads to spreading. Colors represent the local integrated signal, 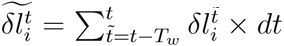, and arrows point towards higher local signal. E) The scaled base contact area *A*(*t*)/*A*(0) as a function of time, with the probability distribution function of vertical displacement shows a net negative response, i.e. bees move upwards on average, similar to experimental observations (see Fig. 2E.

As expected, we see that for the same forcing, the maximum amplitude of the local strains increases as the cluster becomes more elongated (Fig. 3A,B, Movie S6). Therefore, these local strains can serve as a signal for the bees to move, and a natural hypothesis is that once the signal is above a certain critical value, the bees move. But how might they chose a direction ? While it may be plausible for the bees to simply move upwards against gravity, it is likely difficult to sense a static force (i.e., gravity) when experiencing large dynamic forcing (i.e., shaking) in a tightly packed assemblage. Instead, we turn to ask whether there are any local signals that would give honeybees a sense of direction. For all clusters, the strains are largest near the base (Fig. 3A,B, Movie S6) and decrease away from it, but in addition, as the cluster becomes more elongated, there are large local strains along the contact line where *x* = ±*L*_1_/2 where the bees are in contact with the baseboard. This is due to the effect of the pendular mode of deformation that leads to rotation-induced stretching in these regions. To quantify how the normal and shear strain vary as a function of the distance from the base, *Z*, we average *δl*_*i*_ and *δθ*_*i*_ over all bees that were at a certain *Z* position at *t* = 0 and define the following mean quantities: *δl*(*Z*) = 〈*δl*_*i*_〉, and *δθ*(*Z*) = 〈*δθ*_*i*_〉, where the bracket <> indicate average over all spring connection at the vertical position 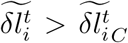. Similar to the experimental data, the simulations show that the displacements *U*_*i*_ for horizontal shaking of elongated clusters are larger in comparison to flattened clusters. As both strains *δl*(*Z*) and *δθ*(*Z*) are largest near the base, *z* = 0 (Fig. 3C, Movie S6), and decrease away from the supporting baseboard, they may serve as local signals that bees at the tip of the cluster respond to by moving up the strain-gradient (Fig. S3–5, Movie S7–8).

This passive signature of a horizontally-shaken assemblage suggests a simple behavioral hypothesis: bees can sense the local variations in the normal strain above a critical threshold, and move slowly up gradients collectively. We note that mechanical strain is invariant to translation and rotation of the whole assemblage, i.e. it is independent of the origin and orientation of the frame of reference, and thus a natural choice. This behavior will naturally lead to spreading of the cluster and thence smaller strains on the cluster. Noting that time scale of the response of the bees is of the order of minutes while the duration of a single period is seconds, it is natural to consider the integrated local normal strain signal: 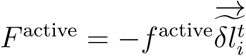 where *T*_*w*_ is chosen to be the period of the shaking (see detailed description in the SI Sec. C). Then our behavioral hypothesis is that when 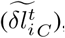 the bee becomes active, and moves in the direction of the time-integrated negative normal strain gradient (i.e. the active force is directed toward a higher local normal strain) according to the simple proportional rule 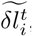 We note that moving up a gradient in time-integrated shear strain would also suffice to explain the observed mechanical adaptation.

We carry out our simulations of the active cluster in two dimensionxs for simplicity and speed (we do not expect any changes in 3D), allowing bonds to break and reform based on proximity similar to how bees form connections, and follow the shape of the cluster while it is shaken horizontally. We find that over time, the cluster spreads out to form a flattened cone (Fig. 3D,E, Movie S7), confirming that the local behavioral rule that integrates relative displacements that arise due to long-range passive coupling in the mechanical assemblage wherein bees actively move up the local gradient in normal strain *δl*_*i*_ does explain our observations.

If sufficiently large dynamic normal strain gradients drive shape adaptation, different shaking protocols that result in lower local strains should limit adaptation. One way is to shake the cluster gently, and this indeed does not drive adaptation (Fig. 2B responding to 0.01g). Another is to shake the cluster in a vertical direction, exciting the spring-like mode of the assemblage. For the same range of amplitudes and frequencies as used for horizontal shaking, our simulations of a passive assemblage show that vertical shaking results results in particles being collectively displaced up and down, with little normal strain. As expected, even in active clusters with the behavioral rule implemented, little or no adaptation occurs as the threshold normal strain gradient is not achieved (Fig. S5–6 and Movie S8). To test this experimentally, we shake the cluster vertically. We see that in this case the cluster shape remains approximately constant (Fig. 4A,B) until a critical acceleration is reached - at which time a propagating crack results in the detachment of the cluster from the board (Movie S9). Quantifying the displacements at the tip for vertical shaking and horizontal shaking are in agreement with our hypothesis that differential normal strain gradients drive adaptation (Fig. 4C, Movie S10).

**Figure 4:**
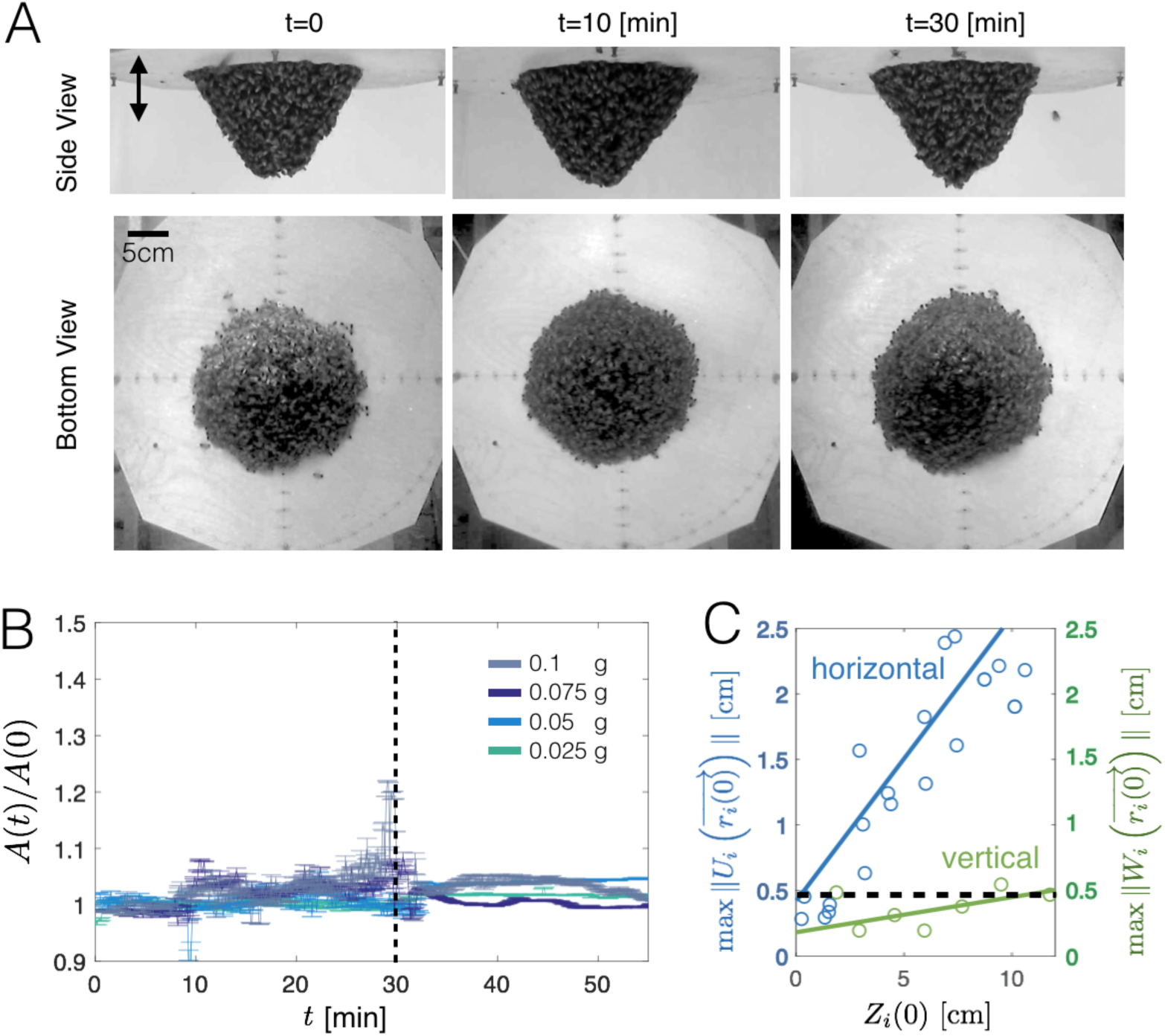
Response to Vertical Shaking. A) Vertical shaking (maximum acceleration 0.05 [g]) of the bee cluster leads to a very small displacement. This is consistent with our simulations (see Fig. S4, SI Sec. D) that vertical shakes do not destabilize the bees differentially. B) Contact area of the base of the cluster relative to its initial area *A*(*t*)/*A*(0) vs. time. Areas are defined as in Fig. 1D. Colors represent results for different accelerations of continuous shaking. C) Maximum displacement at the tip of a tall cluster as a result of a single horizontal and vertical shake. Bees do not respond or change the shape of the cluster when subjected to vertical shaking (green), but do respond substantially when shaken horizontally (blue). Black dotted line represents the experimentally observed threshold value to initiate active behavior.

Our study has shown how dynamic loading of honeybee swarm clusters leads to mechanical adaptation wherein the cluster spreads out in response to repeated shaking. We show that this morphological response increases the mechanical stability of the cluster. A computational model of the bee cluster treated as a passive mechanical assemblage suggests that there are strain gradients that can be used by bees as a local signal to drive movement in the direction of higher strains and causing the cluster to flatten. This behavioral response improves the collective stability of the cluster as a whole via a reversible shape change, at the expense of increasing the average mechanical burden experienced by the individual. Introducing elastic interactions between the bees naturally allows for long-range signaling via physical cues, complementing the more traditional view of collective behavior via stigmergy (*16*) wherein organisms respond to local chemical cues with little or no long range effects. Given the tensorial non-local nature of elastic interactions in assemblages of social insects, perhaps this is just the tip of an iceberg that hides the many ways in which organisms take advantage of physical interactions and simple behavioral rules to adapt to changing mechanical environments.

